# A Novel Approach to Reducing Spinal Implant-Associated Infections

**DOI:** 10.1101/2024.08.30.610521

**Authors:** Mitsuhiro Nishizawa, Diane Hu, Hassan Serhan, Bahram Saleh, Ralph S Marcucio, Kazuhito Morioka

## Abstract

Implant-associated infection is a severe complication of instrumented spinal surgery. Preventive measures against biofilm formation, such as chemical modifications of devices with antibiotics or bactericidal substances, have been explored to decrease infection rates and enhance therapeutic outcomes. However, this approach remains challenging, and the safety issues associated with antibiotic resistance and cellular toxicity need to be addressed. Hence, there is an unmet clinical need for implants that resist biofilm formation. Ultrafine-grained material (UFG), which has a nanoscale grain structure, is a novel form of metal with exceptional mechanical and biocompatibility properties that may also have antimicrobial activity. This study investigates the extent to which UFG stainless steel implants resist biofilm formation by *Staphylococcus aureus in vitro* and *in vivo*. The results show that the UFG stainless steel wire significantly reduced biofilm formation at early time points after bacterial infection compared to a standard wire. In mouse models of implant-associated infection below the skin or the fascia, the UFG wire had significantly less biofilm formation at early time points. The UFG wire appeared to delay initial bacterial adhesion and biofilm maturation. These findings suggest that UFG implants may resist initial biofilm formation and maturation, leading to promising antibacterial materials for patients.

## Introduction

Implant-associated infection is a serious complication following spinal fusion surgery^1^. These conditions are associated with increased morbidity and mortality in affected patients as well as significant costs to the healthcare system^2^. Despite advances in surgical techniques including increased sterility levels, prophylactic antibiotic therapy, and intensive perioperative care, the incidence of implant-associated infection following adult spinal instrumentation varies from 0.7% to 20%^3^. As a result of the increasing demand for spinal surgeries to improve the patient’s quality of life due to the growth of an aging population, implant-associated infections will become a growing burden on healthcare systems in the coming decades^4,5^.

Bacterial adhesion and subsequent biofilm formation on implant surfaces cause implant-associated infection^6-9^. Microorganisms that form biofilms are able to evade the host immune system. Bacterial biofilms are sessile microbial communities that exhibit changing patterns of growth, gene expression, and protein production^10-12^. The biofilm life cycle consists of four stages: adhesion, microcolony formation, maturation, and dispersion. Biofilm formation is triggered by adhesion of planktonic cells to implant surfaces, leading to microcolony formation through cellular aggregation via intercellular adhesion. Subsequent cellular aggregation and multilayered macrocolony formation lead to maturation, with a mix of live and dead bacteria^12^. Finally, planktonic cells are dispersed from fully mature biofilms. Thus, minimizing bacterial adhesion and preventing initial phases of biofilm formation may be crucial strategies to lower the risk of implant-associated infection.

Research and development of antibacterial implants could significantly advance surgical procedures to address these clinically intractable problems. Various surface modifications have been tested to prevent the initial adhesion of bacteria, including antibiotics, low-molecular-weight organic compounds, metal ions, and nanoparticles^13-16^. Among these, antibiotic-coated implants have not been widely used due to adverse reactions, such as cell toxicity, allergies, and altered bacterial flora, leading to antibiotic-resistant strains^13^. Therefore, there is still an unmet clinical need for antibacterial implants that balance efficacy, stability, durability, and safety to reduce the risk of implant-associated infection.

Ultrafine-grained (UFG) materials are state-of-the-art metals and alloys, having advanced multifunctional properties that can be applied to medical devices, such as orthopaedic implants^17^. With nanostructured grain sizes ranging between 100 and 1000 nm, UFG materials exhibit superior mechanical properties such as enhanced strength and fatigue resistance^18^. These characteristics, along with their efficient fabrication and simple processing procedures, make UFG materials highly suitable for a wide range of industrial applications. Furthermore, UFG materials are highly biocompatible and capable of effectively reducing the inflammatory response of the host immune system, making them excellent candidates for clinical applications^17,19,20^. Grain size reduction to nanometer levels can generally affect the interaction of the surface with tissues and cells. These materials may facilitate migration, adhesion, and proliferation of different cell types, such as mesenchymal stem cells, osteoblasts, and fibroblasts^21-24^. While the precise mechanisms underlying these favorable responses are not known, UFG materials typically possess greater surface energy and hydrophilicity compared to conventional coarse-grained materials, leading to enhanced protein adsorption, cell adhesion, and cellular interactions^24,25^. Meanwhile, the effect of UFG materials on bacteria remains controversial. Recent studies show that UFG titanium implants may promote bacterial adhesion^28,49^. However, nanostructured materials reduce bacterial adhesion and proliferation on the material surface^23,26,27^. Bagherifard et al. found that the UFG stainless steel implant reduced the adhesion of gram-positive bacteria^23^, and Yu et al. showed that the UFG stainless steel implant had a lower amount of bacteria on its surface compared to coarse-grained austenitic stainless steel^27^. Hence, UFG, depending on the type of metal used, may be a promising material for biocompatible antimicrobial implants. However, in order to demonstrate the antibacterial properties of UFG materials, it is necessary to evaluate their *in vivo* efficacy, which has not been examined yet.

In this study, we assessed the impact of UFG stainless steel implants on the biofilm formation of *Staphylococcus aureus in vitro* and *in vivo*. Since implant-associated infections following instrumented spinal surgery can spread to either subcutaneous or subfascial space^28^, we examined the antibacterial effects in two different *in vivo* conditions: 1) implants were placed into a pre-existing air pouch located between the subcutaneous tissue and paraspinal fascia, and subsequently inoculated with, and 2) implants were inserted into the lumbar spinous process, and subsequently inoculated with bacteria between the paraspinal fascia and vertebrae as a spinal implant infection model. We hypothesized that UFG stainless steel implants would significantly retard biofilm formation both *in vitro* and *in vivo*. These investigations provide a substantial contribution to the examination of the material potential as well as a better understanding of the mechanisms of antibacterial activity that may promote resistance to implant-associated infection.

## Results

### In vitro antibacterial effect of ultrafine-grained stainless steel implant

For this work, we used *Staphylococcus aureus* Xen36 to investigate the *in vitro* antibacterial properties of ultrafine-grained (UFG) stainless steel implants. We evaluated the biofilm that formed on the wire surfaces at 4 hours, 1 day, 3 days, or 7 days of incubation. The crystal violet (CV) -stained biofilm became visible after 3 days of incubation and subsequently increased over time on both types of wires (Fig. 1A). The absorbance of CV-stained biofilms showed no significant differences between the two types of wires at any time points (Fig. 1B). To quantify the overall amount of biofilm on the wire surfaces, we used the quantitative polymerase chain reaction (qPCR) analysis to determine the relative expression levels of *luxA* and 16S rRNA. Xen 36 is a bioluminescent strain of *Staphylococcus aureus* containing the *luxABCDE* operon^32^. The intensity of the luminescence signal has demonstrated a reliable correlation with the total number of bacteria within biofilms in recent studies^32-35^, indicating that measuring the overall amount of biofilm, which includes both live and dead bacteria, is feasible by quantifying expression levels of the lux gene. The 16S rRNA, which is a component of the prokaryotic ribosomal 30S subunit found in all bacterial genomes, is widely used to quantify the overall amount of biofilm because of its stable expression levels^36^. Our results showed that there was no significant difference in the expression of either gene between both types of wires (Fig. 1C). To determine the total number of live bacteria within the biofilm on the wire surfaces, we performed a colony-forming unit (CFU) assay. We observed that the UFG stainless steel wire had significantly fewer live bacteria than the standard wire at 4 hours (p = 0.0005), 1 day (p = 0.0001), and 3 days (p = 0.0314) after incubation (Fig. 1D). The early stages of biofilm formation by Staphylococcus aureus can be identified by expression of intercellular adhesion *(icaA)* and the clumping factor A (*clfA)*. To assess the early biofilm on the wire surfaces, the expression levels of *icaA* and *clfA* were determined by qPCR. Our results showed that the UFG stainless steel wire had significantly lower expression of *icaA* than the standard wire at 1 day (p = 0.042) and 3 days (p = 0.004) of incubation (Fig. 1E). Furthermore, the UFG stainless steel wire revealed a significant decrease in the expression of *clfA* compared to the standard wire at 3 days of incubation (p = 0.007).

**Fig 1.**
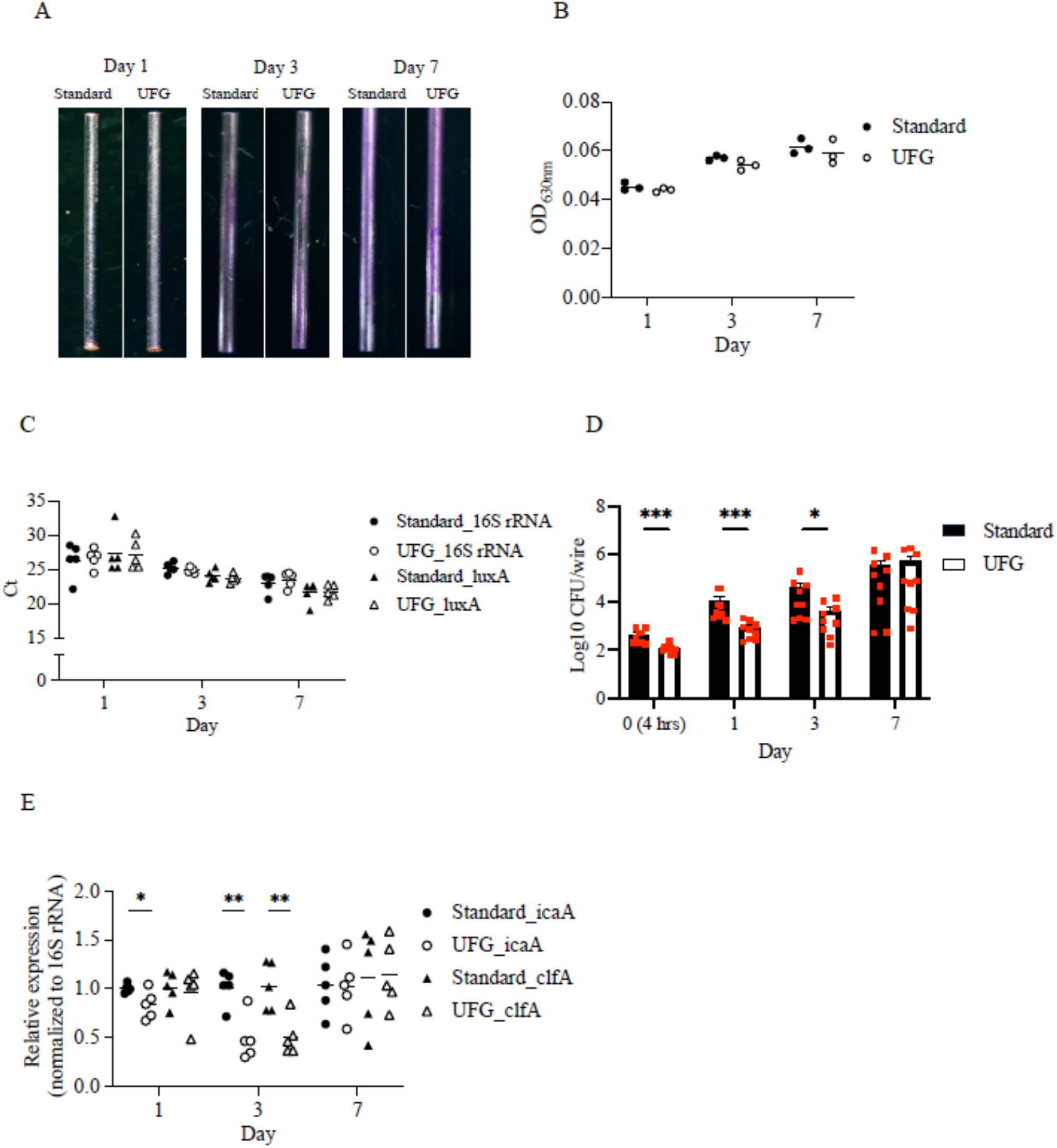
*In vitro* effects of ultrafine-grained stainless steel implants on *Staphylococcus aureus* biofilms. (**A**) The representative images of crystal violet-stained biofilms formed by *Staphylococcus aureus* on the surface of the ultrafine-grained (UFG) stainless steel wire and the standard wire at one day (left), three days (middle), and seven days (right) of incubation. (**B**) The absorbance measurement of crystal violet-stained biofilms on the wire surfaces at the indicated time points (n = 3 per wire type). (**C**) The gene expression of the lux operon (*luxA*) and 16S rRNA were used to measure the overall amount of biofilms on the wire surfaces at the indicated time points (n = 5 per wire type). The cycle threshold (Ct) values are inversely correlated with the quantity of target nucleic acid and bacterial load. (**D**) The colony-forming unit (CFU) assay was employed to quantify the total number of live bacteria present within the biofilms on the wire surfaces at the indicated time points (n = 9 per wire type). (**E**) The relative expression of the intercellular adhesion (*icaA*) gene and the clumping factor A (*clfA*) gene were measured to assess the early development of biofilms on the wire surfaces (n = 5 per wire type). The horizontal line on the dot plot represents the mean values. The dots indicate individual data points. The bar graph represents the mean values ± standard error of the mean. Statistical significance was tested by one-way ANOVA (**E**), or two-sided Mann–Whitney U test (**D**). ^*^p < 0.05, ^**^p < 0.01, ^***^p <0.001.

### In vivo antibacterial effect of ultrafine-grained stainless steel implant in subcutaneous pouch infection model

For *in vivo* assessments, we initially evaluated the effects of UFG stainless steel implants on biofilm formation using a subcutaneous pouch infection model. In general, *in vitro* biofilm formation is highly dependent on the conditioned medium in the absence of the immune system^29^, and the direct comparison between *in vitro* and *in vivo* results is challenging^30^. Our experimental model was created to mimic the clinical characteristics of implant-associated infection following instrumented spinal surgery, such as the development of an abscess around implants beneath the skin. By integrating two types of wires and inserting them as a single unit into the infection site, the subcutaneous pouch infection model allows for the evaluation of biofilms formed on each surface under the same infection conditions within the same animal.

Both wires were placed into the enclosed subcutaneous space containing Xen36. Following incubation, a fully developed abscess encasing both wires formed in the subcutaneous region (Fig. 2A). The CV-stained biofilms were detected 1 day after inoculation (Fig. 2B) and were not different between the wires at any time point (Fig. 2C). However, the UFG stainless steel wire had significantly fewer live bacteria than the standard wire 1 day after inoculation (p = 0.011; Fig. 2D). Additionally, the UFG stainless steel wire had a significantly lower expression of *icaA* than the standard wire at 1 day after inoculation (p = 0.002) (Fig. 2E).

**Fig 2.**
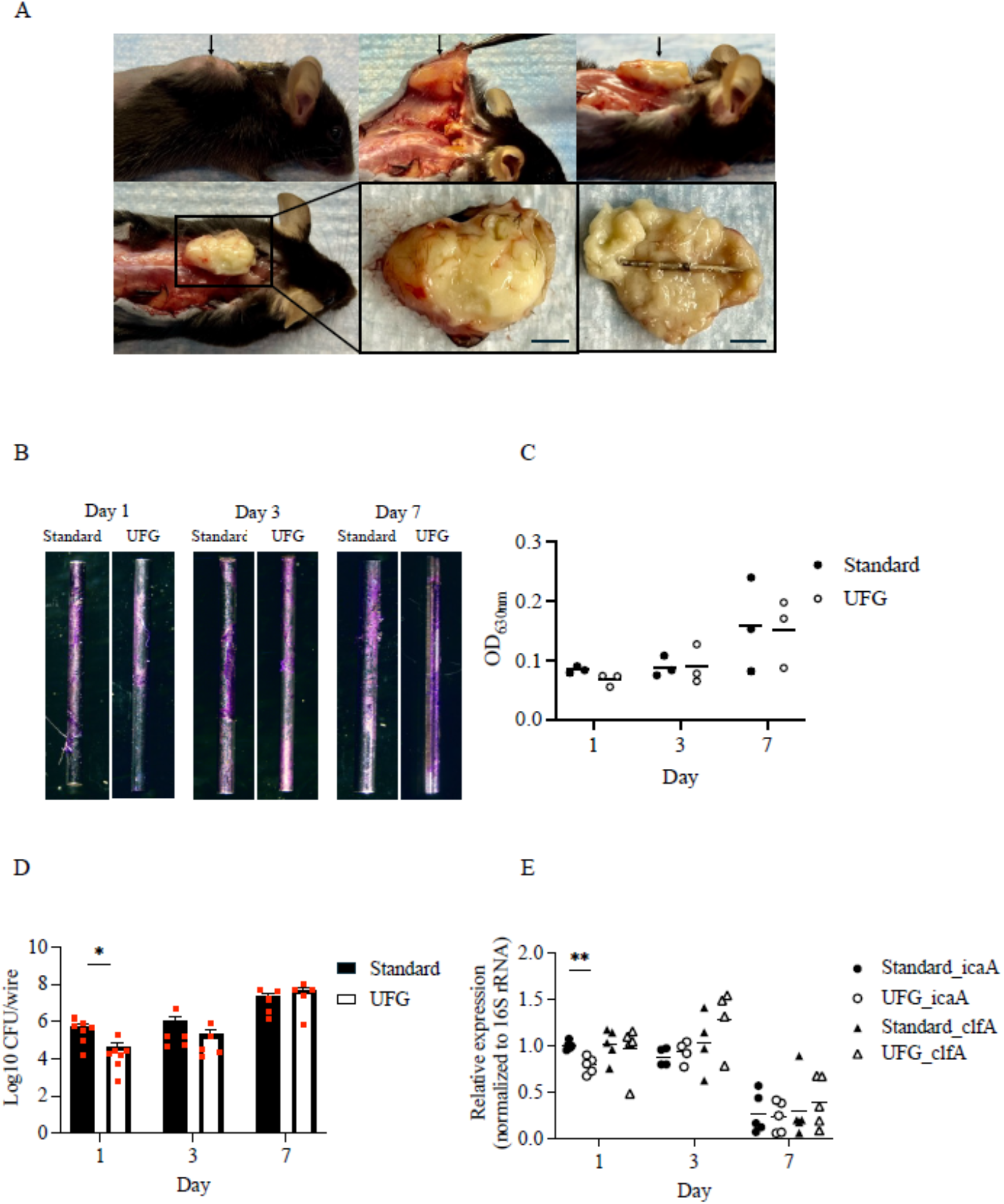
*In vivo* effects of ultrafine-grained stainless steel implants on *Staphylococcus aureus* biofilms in subcutaneous pouch infection. (**A**) The representative image of the subcutaneous pouch infection model at seven days after inoculation with *Staphylococcus aureus* (upper left). The well-developed abscess was located beneath the skin (upper middle) and above the paraspinal fascia (upper right). The wire combining the ultrafine-grained (UFG) stainless steel wire and the standard wire was present within the solitary subcutaneous abscess (lower images). The arrow indicates a subcutaneous abscess. Scale bars, 5 mm. (**B**) The representative images of crystal violet-stained biofilms on the wire surfaces at one day (left), three days (middle), and seven days (right) after inoculation. (**C**) The absorbance of crystal violet-stained biofilms was measured to assess the overall amount of biofilms on the wire surfaces at the indicated time points (n = 3 per wire type). (**D**) The colony-forming unit (CFU) assay was used to measure the total number of live bacteria within the biofilms on the wire surfaces at one day (n = 7 per wire type), three days (n = 5 per wire type), and seven days (n = 5 per wire type) after inoculation. (**E**) The gene expression of intercellular adhesion (*icaA*) and clumping factor A (*clfA*) was quantified to evaluate the early development of biofilms on the wire surfaces at one day (n = 5 per wire type), three days (n = 4 per wire type), and seven days (n = 5 per wire type) after inoculation. The dots represent individual data points, and the mean values ± standard error of the mean are shown in the bar graph. Statistical significance was tested by one-way ANOVA (**E**), or two-sided Mann–Whitney U test (**D**). ^*^p < 0.05, ^**^p < 0.01.

### In vivo Antibacterial effect of ultrafine-grained material in spinal implant infection model

Next, we evaluated the antibacterial effects of UFG stainless steel implants on biofilm formation using a lumbar spinal infection model. To evaluate the biofilm formed on two types of implants in the same individual and under the same infection conditions, we modified the well-established mouse model of spinal implant infection^31,32^. By adequately detaching the soft tissues from both sides of the vertebrae, L-shaped wires were inserted bilaterally into the spinous process. After inoculation with Xen36 into the right and left subfascial spaces, an extensively formed subfascial abscess developed in the midline of the lumbar spine over time, but both wires were not completely encased in the abscess (Fig. 3A). The presence biofilms were observed beginning on 1 day after inoculation (Fig. 3B) and were not different between wires at any time point (Fig. 3C). However, again, the UFG stainless steel wire had significantly fewer live bacteria than the standard wire at 3 days (p = 0.015) (Fig. 3D). No differences were found in the expression of *icaA* and *clfA* between the wires at any time points (Fig. 3E).

**Fig 3.**
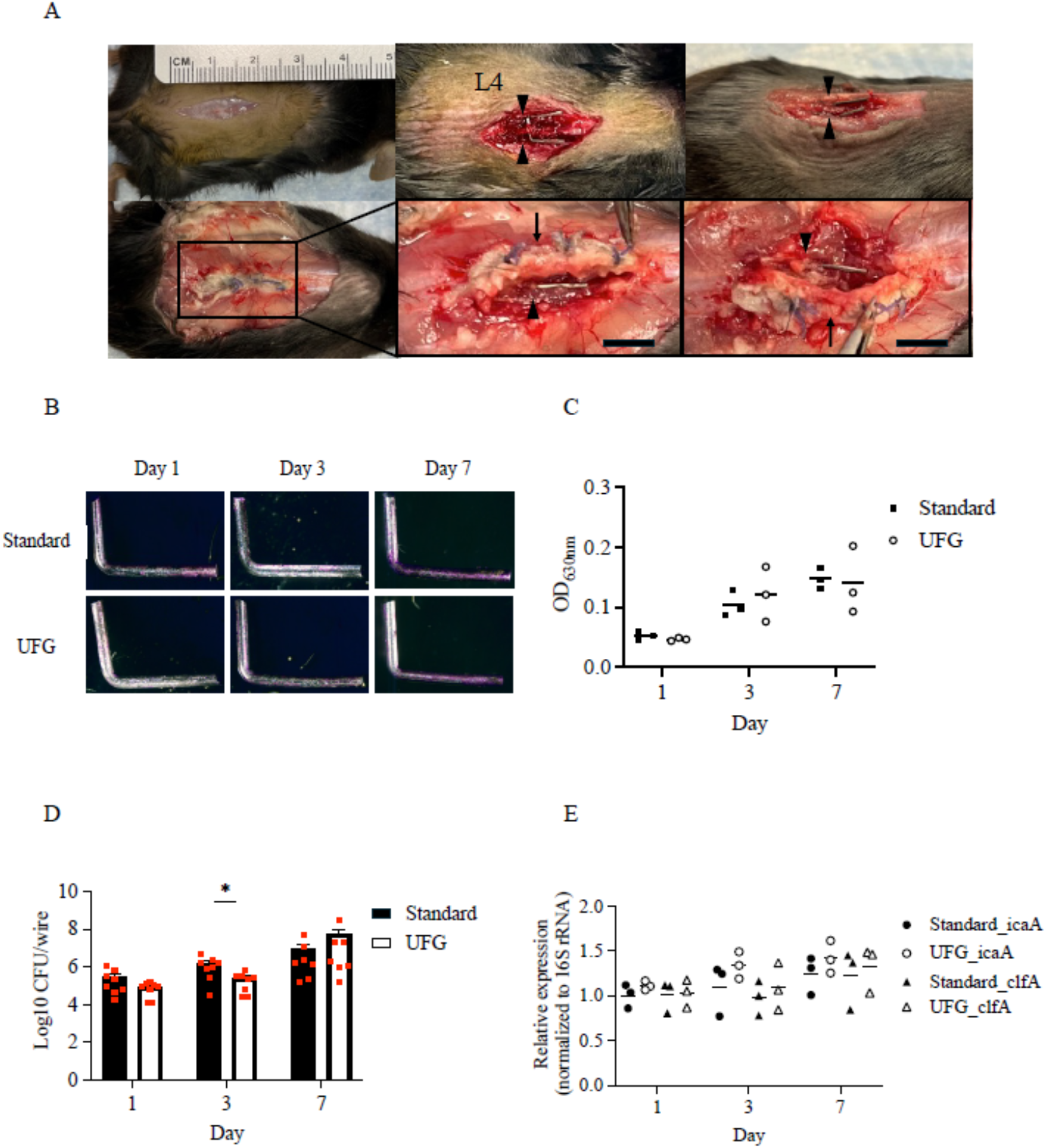
*In vivo* effects of ultrafine-grained stainless steel implants on *Staphylococcus aureus* biofilms in spinal implant infection. (**A**) The representative images of the spinal implant infection model. To expose the spinous process of the fourth lumbar vertebra (L4), a skin incision was made in the middle of the lumbar spine (upper left). After two holes were created in the spinous process, an L-shaped ultrafine-grained (UFG) stainless steel wire or an L-shaped standard wire was inserted into the hole from either the left or right side (upper middle, right). The extensively formed abscess was located at the paraspinal fascia (lower left) and above the vertebrae (lower middle, right) at seven days after inoculation with *Staphylococcus aureus*. Both lateral wires were present adjacent to the abscess (lower middle, right). The arrow denotes a subfascial abscess, and the arrowhead indicates a wire. Scale bars, 5 mm. (**B**) The representative images of crystal violet-stained biofilms on the surfaces of the L-shaped wire at one day (left), three days (middle), and seven days (right) after inoculation. (**C**) By measuring the absorbance of crystal violet-stained biofilms, the overall amount of biofilms on the wire surfaces was evaluated at the indicated time points (n = 3 per wire type). (**D**) The total number of live bacteria within the biofilms on the wire surfaces was quantified by the colony-forming unit (CFU) assay at one day (n = 8 per wire type), three days (n = 8 per wire type), and seven days (n = 7 per wire type) after inoculation. (**E**) The gene expression levels of intercellular adhesion (*icaA*) and clumping factor A (*clfA*) were measured to assess the early development of biofilms on the wire surfaces at the indicated time points (n = 3 per wire type). The dots in the graph indicate individual data points, while the bar graph displays the mean values ± standard error of the mean. Statistical significance was tested by two-sided Mann–Whitney U test (**D**). ^*^p < 0.05.

## Discussion

In this study, we found that UFG stainless steel materials have the potential to resist the early stages of biofilm formation when exposed to high amounts of *Staphylococcus aureus*, even in the absence of antibiotics. Our *in vitro* results showed a significant reduction in live bacteria within the biofilm on the surface of the UFG stainless steel implant up to 3 days of incubation in the 3 different conditions of implant-associated infection. Considering that biofilm formation occurs rapidly^33^, delays in the biofilm formation on the UFG stainless steel wires may confer an advantage by delaying this process. Our *in vivo* data are consistent with the findings from the *in vitro* results, indicating that UFG stainless steel implants may inhibit proliferation of viable bacteria within the biofilm. Furthermore, we found a significant decrease in the expression levels of the *icaA* or *clfA* genes, encoding the proteins associated with the extracellular matrix that plays a crucial role in the early stages of biofilm formation^7,9-12,34^. In the spinal infection model, no statistical difference in gene expression was observed between the two types of wires, and there was no notable alternation in gene expression over time, which may be attributable to the fact that the wires were not completely encased in the abscess. During the life cycle of the biofilm, the initial adhesion of bacteria to the material surface is regulated by the interactions with the material surface through electrostatic, hydrophobic, and Van der Waals forces. Subsequently, irreversible adhesion requires cell wall-anchored proteins encoded by the *clfA* gene that binds to host fibrinogen to facilitate microcolony formation. The polysaccharide intercellular adhesion protein (PIA), encoded by the *icaADBC* locus, plays a major role in cell-cell adhesion and promotes biofilm maturation^35-37^. These results may indicate that UFG stainless steel implants may resist initial bacterial adhesion and subsequent biofilm development.

The delay in establishment of the biofilm may have significant clinical value. The lower levels of contaminating bacteria in a clinical setting may have less chance of establishing a biofilm on the implant in the presence of prophylactic antibiotics and the immune response. *In vitro*, there were fewer live bacteria on the UFG wires at early time points. This was accompanied by the decreased by the expression of *icaA* and *clfA*.

*In vivo*, we observed the differences in the amount of live bacteria on the UFG wires. In the pouch model, fewer live bacteria were present at day 1, which is in line with our *in vitro* data. However, in the spine model, we observed fewer bacteria on the UFG wires at day 3. The differences of these results could be attributed to several factors, such as the location of the infection site (subcutaneous versus subfascial) and the volume of bacterial inoculation (large versus small). Moreover, the two *in vivo* models differ in the spatial relationship between the wire and the abscess, which may also influence the results. In the pouch model, the wire is completely surrounded by the abscess, whereas in the spine model, the abscess extends primarily dorsal to the wire.

This study has several limitations. First, the investigation was conducted using only one strain of *Staphylococcus aureus*, Xen36, and a single type of UFG material, stainless steel UFG with a grain size of 500 nm, which is produced by the warm oval square rolling process. The bacterial cell behavior and the subsequent inflammatory response can vary depending on the grain size, nanoscale topographic roughness, and the size and shape of the implant surface, the bacterial species, and even the specific process to create the UFG materials^23-27^. Further research is required to fully understand these interactions. Secondly, this study did not concurrently evaluate the mechanical properties of the metal used, such as roughness, surface charge, and wettability, alongside biofilm assessment. Thirdly, this study only compared biofilm formation at relatively early time points, up to 7 days post-infection. Therefore, the effects of UFG materials on bacteria at later time points remain unclear, and evaluating these later stages is crucial for considering the application of UFG materials in orthopedic implants. Fourth, we used very large bacterial inoculums which may not reflect a clinical scenario, and we did not provide prophylactic antibiotics or treat the infections as would be standard of care.

In conclusion, to our knowledge, this is the first *in vivo* study to demonstrate that UFG stainless steel possesses potential antibacterial properties capable of resisting the adhesion of bacteria and subsequently microcolony formation and biofilm maturation. The unique characteristics of UFG materials, produced without altering their chemical composition, address concerns of cytotoxicity often associated with other antibacterial technologies. Therefore, incorporating UFG materials in the manufacture of orthopedic implants or medical devices could uniquely balance antibacterial properties with biocompatibility, making them highly suitable for these applications.

## Methods

### Bacteria Preparation

Xen 36 is a strain of *Staphylococcus aureus* that exhibits bioluminescence, which was acquired from the American Type Culture Collection (ATCC49525) (PerkinElmer, Inc., Waltham, MA). This strain harbors a genetically modified luxABCDE operon and a gene conferring resistance to kanamycin. These features enable us to precisely identify these bacteria using qPCR to target the genes in the lux operon and to cultivate uncontaminated cultures of these bacteria in the presence of kanamycin. Xen36 was streaked on tryptic soy agar (TSA) plates containing tryptic soy broth (TSB) and 1.5% mannitol salt agar. The plates were then incubated at 37°C overnight. A single colony was selected and cultured overnight in a TSB medium with 200 µg/ml kanamycin to eliminate any contaminations at 37°C in a shaking incubator (200 rpm). The bacterial solutions were resuspended in TSB at a concentration of 1.0 × 108 colony-forming unit (CFU)/ml, determined by measuring the absorbance at 630 nm using a microplate reader (800TS, Agilent Technologies, Inc., Santa Clara, CA).

### Implant

Ultrafine-grained (UFG) wires manufactured of stainless steel (SUS316L), 8 mm in length and 0.5 mm in diameter, were utilized for both *in vitro* and *in vivo* experiments. Unprocessed SUS316L wires were used as the standard wire to compare the impacts of the UFG wires. This particular austenitic stainless steel is frequently used in various industries, including biomedical applications. A severe plastic deformation process was used to produce UFG stainless steel with a grain size of 500 nm from commercially available SUS316L, which initially had a grain size of over 10 µm (Komatsuseiki Kosakusho Co., Ltd., Japan). The grain size was determined using the Electron Back Scattered Diffraction Pattern detector (EBSD DigiView, EDAX, LLC, Pleasanton, CA). All wires and supplies underwent autoclaving prior to the studies.

### In vitro Biofilm Formation

The wires were placed into the wells of a 24-well plate containing 1 ml of a 1.0 × 10^8^ CFU/ml solution for Xen36 culture. The plates were incubated at 37°C with shaking at 200 rpm. The wires were rinsed with deionized (DI) water three times, and TSB containing 200 µg/ml kanamycin was replaced every 24 hours. The wires were extracted from the medium at 4 hours, one day, three days, and seven days of incubation. At each time point, the wires were gently rinsed with DI water to eliminate any germs that were loosely attached to the surface. The biofilm on the wires was evaluated by the CV analysis, the qPCR analysis, and the CFU assay as described below. All experiments were performed in triplicate.

### Animals

All animal procedures were approved by the Institutional Animal Care and Use Committee (IACUC) at the University of California, San Francisco (UCSF). The use of bacteria was performed in a BSL2 facility after consultation with and approval by the UCSF Biosafety Hazard Program, administered by UCSF Environmental Health and Safety. We have used biosafety level 2 (BSL2) pathogens in the animal facility. Adult C57/B6 mice were used in *in vivo* experiments. To minimize pain, mice were anesthetized during all surgical procedures. The mice were anesthetized using inhalation of 2% isoflurane. The mice were carefully monitored after surgery for signs of discomfort, pain, or distress and administered analgesics (buprenorphine; 0.1 mg/kg) as appropriate throughout the course of the experiments or euthanized per the recommendations of the UCSF veterinary consultants. To minimize distress, mice were housed communally and provided environmental enrichment devices as allowed by Federal Regulations and our IACUC protocols. Routine care for animals was provided by the UCSF Laboratory Animal Resource Center. The euthanasia was conducted in accordance with the American Veterinary Medical Association (AVMA) Guidelines for the Euthanasia of Animals.

### Mouse Model of Subcutaneous Pouch Infection

Seven days prior to the induction of bacterial infection, the following procedure was implemented to generate the air pouch in mice: Mice were anesthetized using inhalation of 2% isoflurane. The dorsal fur from the sacrum to the upper thoracic spine was removed with clippers. Appropriate depth of anesthesia was confirmed by ensuring that respiration remained rhythmic, slower than when awake, and unresponsive to noxious stimuli. The base of the mouse’s neck was then gently pinched and lifted upward to create space between the subcutaneous tissue and the fascia. A 27-gauge needle was used to inject 3 ml of sterile air subcutaneously into this space, thereby forming the air pouch. To maintain the cavity and mature the pouch, 3 ml of air was reinjected every 2 days. Before each inflation, the air was aspirated from the pouch to confirm the proper placement of the needle tip. Seven days after the initial air injection, the mouse was anesthetized, and appropriate depth of anesthesia was confirmed as previously described. The skin was sterilized with isopropyl alcohol and betadine. Buprenorphine (0.1 mg/kg) was injected subcutaneously just prior to the surgery, and a 5 mm midline longitudinal incision was made at the proximal end of the pouch. An implant, which included two types of wires connected vertically using a 20 µl tip cut to a length of approximately 3 mm, was gently inserted into the pouch. 3 ml of 1.0 × 10^5^ CFU/ml S. aureus Xen36 culture was injected into the pouch, and the skin was closed with a wound clip and sealed with topical skin adhesive. The mice were placed on a heating pad and monitored for return to normal activity. At 1-, 3-, and 7-days post-infection, the mice were euthanized with carbon dioxide inhalation, the samples were harvested, and the biofilm on the implants was evaluated by CV assay, CFU counting, and qPCR as described below.

### Mouse Model of Postoperative Spinal Implant Infection

Mice were anesthetized (inhalation 2% isoflurane) and the hair on the dorsal area from the sacrum to the upper thoracic spine region was shaved with clippers. The skin was sterilized with isopropyl alcohol and betadine solution. After confirming the appropriate depth of anesthesia, a longitudinal 3 cm incision was made at the dorsal aspect of the centered lumbar spine region. By cutting the fascia along the midline, the spinous processes were exposed, and paraspinal muscles were peeled away from the spine bilaterally. Using a 25G needle, two holes were created in the spinous process. Subsequently, wires bent into an L-shape measuring 3 mm x 5 mm were employed, with the shorter arm of the pin being passed through each of these holes from the left and right sides, respectively, to anchor the implants to the spine, and the long arm extended toward the head of the mouse^32^. An inoculation of 1 × 10^3^ CFU (10 µl of 1.0 × 10^5 CFU/ml) of Xen36 was applied in the cavity to ensure all solution contacted the implants. The fascia layer was sutured watertight with 4-0 Vicryl, and the skin was closed using clips. Buprenorphine (0.1 mg/kg) was injected subcutaneously, and the mice were placed on a heating pad and monitored for return to normal activity. At 1-, 3-, and 7-days post-infection, the mice were euthanized with carbon dioxide inhalation, the samples were harvested, and the biofilm on the implants was evaluated by CV assay, CFU counting, and qPCR as described below.

### Crystal Violet Assay

The biofilm on the implant surface was stained with CV staining and visualized with a compound light microscope^38^. Wires were gently rinsed with DI water to remove any loosely adherent bacteria from the surface. The biofilms on the wires were fixed using 100% ethanol for one minute and then left to air dry for three minutes. Subsequently, the fixed biofilms were stained with a 0.1% CV reagent for fifteen minutes. The stained specimens were washed with DI water and imaged using a light microscope. Then, the wires were placed in wells of 96-well plates containing 250 µl of 33% acetic acid to solubilize the CV staining. 200 µl of the suspension was then transferred to new wells, and the absorbance was measured at 630 nm using a microplate reader. All measurements were performed in triplicate.

### Determination of Viable Bacteria Cell Number by CFU Counting

To quantify the number of living bacteria in the biofilm on the implants, we determined CFU^38^. Wires were gently rinsed with DI water to remove any loosely adherent bacteria from the surface and transferred to a 1.5 ml tube containing 200 µl of 10x trypsin^39^. After incubation at 37°C for 1 hour, the wires were vortexed for 1 minute and then sonicated for 10 minutes in a water bath with an output of 100 W. This was then followed by an additional vortexing step for 30 seconds to detach the biofilms from the wire surfaces and disperse them into the suspension. Each supernatant was serially diluted 10-fold, and aliquots were plated onto TSA plates in triplicate. The colony counts were determined after an overnight incubation at 37°C.

### Quantitative Polymerase Chain Reaction

We used qPCR analysis for the 16sRNA and lux operon (*luxA*) to quantify bacterial load on the surface of the implants ^40,41^. Each pin was serially dipped three times in wells containing 1 ml of PBS to remove planktonic microorganisms and transferred to a 1.5 ml tube containing 600 µl of TSB media. The plasmid DNA was extracted from the bacteria using the Zyppy Plasmid Miniprep Kit (ZYMO RESEARCH). The final sample of DNA was eluted in 30 µl of the elution buffer, and the concentration of collected plasmid DNA was measured using NanoDrop (Thermo Fisher Scientific). Quantification was performed using a Light Cycler instrument, and the threshold cycle (Ct) was determined. qPCR analysis for the 16sRNA and *luxA* was used to quantify bacterial load on the surface of the wires. PCR amplification was performed in a total reaction mixture volume of 10 µl. The reaction mixture contained 5 µl SYBR Green, 1.0 µl of forward and reverse primer, 3 µl of 1:10 diluted plasmid DNA obtained from the sample, and 1 µl of nuclease-free water. The primers used were as follows: for *16sRNA*, 5’-GTGGAGGGTCATTGGAAACT-3’, and 5’-CACTGGTGTTCCTCCATATCTC-3’, for *luxA*, 5’-GAGCATCATTTCACGGAGTTTG-3’, 5’-ATAGCGGCAGTTCCTACATTC-3’. The thermal cycling protocol was as follows: initial denaturation for 2 minutes at 94°C, followed by 40 cycles of 15 seconds at 94°C, 30 seconds at 60°C, and 30 seconds at 72°C. The fluorescence signal was measured at the end of each extension step at 72°C. We generated a calibration curve, created with plasmid DNA purified directly from an overnight pure Xen36 culture^42^. The mean of the three-cycle threshold (Ct) values was plotted against a calibration curve to estimate the bacterial load attached to the implant. To further investigate, we examined the expression levels of genes encoding the extracellular matrix and associated proteins (*icaA* and *clfA*) that play an important role in the process of biofilm initiation and early-stage biofilm maturation. Subsequently, PCR amplification was performed under the same PCR conditions, using 3 ng of DNA from each sample and the primers as follows: for*16sRNA*, 5’-GTGGAGGGTCATTGGAAACT-3’, and 5’-CACTGGTGTTCCTCCATATCTC-3’, for *icaA*,5’-GAGGTAAAGCCAACGCACTC-3’, and 5’-CCTGTAACCGCACCAAGTTT-3’,for *clfA*, 5’-TACAAGTGCGCCTAGAATGA-3’ and 5’-TTTGACATAACCTGCTTGGT-3’. The 16sRNA gene was used as an internal control, and the relative gene expressions of these genes were calculated by the 2-ΔΔCT method.

### Statistical analysis

The statistical analysis was performed using IBM SPSS Statistics for Windows, version 29.0 (IBM Corp., Armonk, NY). All data are presented as mean ± standard error. The correlation between continuous variables was examined using Spearman correlation analysis. The differences between the two types of wires were assessed using one-way analysis of variance (ANOVA) or two-sided Mann–Whitney U test. Statistically significant values were defined as p < 0.05. All bar and line graphs were generated using GraphPad Prism version 10.2.2 software for Mac (GraphPad Software, LLC, San Diego, CA). The statistical significance of the observed differences between the two types of wires was indicated with a bar and an asterisk according to the following probabilities: ^*^p < 0.05, ^**^p < 0.01, ^***^p < 0.001.

## Data availability

The datasets generated and analyzed during the current study are available from the corresponding authors upon reasonable request.

## Acknowledgments

This research was partially funded by an NSF Industry/University Cooperative Research Program called the Center for Disruptive Musculoskeletal Innovations (IIP-1916629). We would like to thank Komatsuseiki Kosakusho Co., Ltd. and Rosies Base, LLC for providing the wires used in our experiments.

## Contributions

HS, BS, RM, and KM contributed to the conception and design of the study. MN and DH performed material preparation, experiments, data collection. MN and KM performed data analysis. MN wrote the original draft of the manuscript. RM and KM revised the manuscript. All authors reviewed and approved the final version of the manuscript.

## Additional information

N/A.

## Competing interests

The authors declare no competing interests.

## Notes

### Competing Interest Statement

The authors have declared no competing interest.

